# Insectivorans and Carnivorans Exhibit the Same Social Brain Relationship as Primates

**DOI:** 10.1101/2025.05.05.652185

**Authors:** R.I.M. Dunbar, Susanne Shultz

## Abstract

It has previously been thought that the social brain hypothesis applies in its full quantitative form only to primates. Using more appropriate brain indices and less error-prone statistical methods, we here show that it does in fact apply to both Insectivorans and Carnivorans, as well as to elephants and orcas. While Insectivorans form a grade on their own, reflecting a simple transition from small-brained solitary social systems to slightly larger-brained pairbonded monogamy, the Carnivorans map directly onto the primate socio-cognitive grades, dividing between loose sociality and more bonded sociality paralleling the pattern observed in monkeys. In addition, the subgroupings of species with multilevel social systems (including hyaenids elephants and orcas) map across the grades in the same way as do primates with similar societies. This suggests that common principles and trajectories underlie social evolution in mammals.

## Introduction

The social brain hypothesis was developed to explain why primates have much larger brains, both absolutely and relative to body size, than all other vertebrate orders [Dunbar 1992]. It argues that the central problem faced by intensely social species is the need to maintain the coherence of groups through space and time in the face of intensely dispersive forces created by the stresses of living in close physical proximity [Dunbar & Shultz 2021a,b, 2023; Shultz & Dunbar 2022; Dunbar 2025]. In primates, this is a consequence of specialised, neurophysiologically expensive cognition associated with managing social relationships that depends principally on the brain’s default mode neural network [Mars et al. 2012; Rushworth et al. 2013] such that the volume of neural material available for key cognitive processing (notably mentalising/perspective-taking and self-control) limits the size of social group that a species can maintain [Dunbar & Shultz 2021a; Dunbar 2025].

Carnivorans comprise a range of families that vary socially from semi-solitary (skunks, racoons, most felids) to pairbonded (most canids) and group-living (some mustelids, hyaenids, lion). Many Carnivorans are highly intelligent, and seem to exhibit at least some of the social and cognitive skills that characterise anthropoid primates [Holecamp et al 2007]. Because their foraging strategies often involve predicting the behaviour of evasive prey, we might expect them to share with anthropoid primates some of the cognitive skills (e.g. perspective taking, mentalising) that underpin primate sociality. Although there have been a number of attempts to test the social brain hypothesis on Carnivorans [Dunbar & Bever 1998; Gittleman 1986; Holecamp et al. 2007; Pérez-Barbería et al. 2007; Shultz & Dunbar 2007, 2010; Finarelli & Flynn 2009; Sakai et al. 2011; Chambers et al. 2021], these have generally concluded that the social brain hypothesis either does not apply to Carnivorans [Finarelli & Flynn 2009; Sakai et al. 2011; Chambers et al. 2021] or does so only in a moderated form (e.g. as a simple binary relationship in which pairbonded species have larger brains than non-pairbonded species [Pérez-Barbería et al. 2007; Shultz & Dunbar 2007]) or only to certain Carnivoran families (e.g. canids but not felids: Shultz & Dunbar 2010]).

One problem that has emerged is that many comparative studies of brain evolution inadvertently introduce unintended statistical errors into their analyses [Dunbar & Shultz 2023]. First, using endocranial volume (ECV) rather than neocortex volume introduces significant measurement error into the brain size variable, resulting in reduced statistical power [Dunbar & Shultz 2021a, 2023]. Although significant results can be obtained using ECV, the fit generally improves, and the slope significantly steepens, if some index of neocortex volume is used to test the hypothesis [Dunbar & Shultz 2021a, 2023]. Second, they invariably ignore the fact that the primate social brain relationship consists of a set of four distinct grades [Dunbar 1993; Dunbar & Shultz 2021a]. In doing so, they fall foul of Simpson’s Paradox (a version of the ecological fallacy), which causes the regression slope to be shallower than it should be, thereby increasing the likelihood of Type II errors [Dunbar & Shultz 2023]. These grades are due to the fact that certain network sizes are attractors for group size because they are more efficient in information transmission [West et al. 2020, 2023]. Dunbar & Shultz [2021a] have shown that, in primates that have multilevel social systems (e.g. chimpanzees, baboons, humans), the characteristic sizes of the different social layers sit on different grades, forming a fractal series. Third, many studies have regressed brain size on group size (especially when using multiple regression), and hence inadvertently tested a radically different hypothesis (namely, does group size constrain brain size, rather than the evolutionarily correct question of whether brain size constrains group size) [Dunbar & Shultz 2023]. The first tests a mechanisms rather than a selection question, and hence often yields weak or non-significant results in contexts where, using the same data, the second yields a strongly significant relationship between group size and brain size [Dunbar & Shultz 2023].

We here follow the approach adopted by [Dunbar & Shultz 2021a] and ask whether neocortex volume (indexed as neocortex ratio) predicts group size in Carnivorans. We extend the analysis to include Insectivorans because this taxon is characterised by forms of sociality (solitary living and monogamy) that are thought to characterise the earliest mammals [Müller & Thalmann 2000]. Since neither of these orders is characterised by primate-like multilevel social systems with a fractal structure (see [Dunbar & Shultz 2021a]), we also include elephants and orcas (which are known to have these kinds of society [Hill et al. 2008]) in order to test whether any non-primates exhibit the same pattern that we find in primates where the units of multilevel societies sit on the grade lines rather than between them (see [Dunbar & Shultz 2021a]). We use neocortex ratio as our metric because, in primates, this consistently yields much cleaner results for both group size and indices of cognition than any other brain index [Dunbar & Shultz 2021a,b, 2023], probably because it indexes relative investment in the functional (‘thinking’) part of the brain (as opposed to that concerned solely with managing somatic tissue).

## Methods

We use data on neocortex and total brain volume for primates and Insectivorans given by [Stephan et al. 1981; Dunbar & Bever 1998] and for Carnivorans by [Röhrs 1986a,b; Röhrs etal.1989; Kamiya & Pirlot 1988a,b; Röhrs, personal communication]. Importantly, these studies all used the same histological methods for estimating brain region volumes. Primate group sizes are from [Dunbar et al. 2018]. Group sizes for Carnivorans and Insectivorans were obtained from a comprehensive search of the primary literature. Group sizes are defined, as in primates, to include immatures when these normally live with the adults. Because, uniquely among the Carnivorans, spotted hyaena (*Crocuta*) live in a form of multilevel society [Holecamp et al. 1997; Smith et al. 2008; Dunbar 2024], defined by a trimodal distribution of group sizes (Fig. S1), we use three group sizes for this taxon: the clan (essentially a matriline with associated males and immatures), a multiclan grouping and a mega-clan grouping [Dunbar 2024]. Group sizes (clique, community, mega-community) and brain volume for chimpanzees are from [Dunbar & Shultz 2021a]; group sizes for elephants (family, bond group, clan) and orca (pod, clan, community) are from [Hill et al. 2008], with brain volumes from [Haug 1970; Hakeem et al. 2005; Wright et al. 2017]. The demands of dolphin echolocation result in an auditory cortex that is 12 times larger than that of humans [Ridgway & Au 2009]; in order to compare like-for-like, orca neocortex volume was reduced by the difference between human and dolphin auditory cortex (calculated from [Rademacher et al. 2001]).

Solitary species are particularly challenging. Many of the primate species once thought to be solitary (e.g. nocturnal prosimians) have, with more intensive field study, turned out to be group-living in that several individuals share a common nest and/or have affiliative social relationships [Nekaris & Bearder 2007]. In effect, early field workers confused foraging groups with social groups [Dunbar & Shultz 2023]. We therefore sought to determine whether or not ‘solitary’ Insectivorans habitually nested together outside of the breeding season (in which case, we counted the nest group size, as in the case of prosimian primates). However, the Insectivorans are extremely poorly studied in the wild. In a few cases (notably the various otter shrew genera), it proved impossible to come to any principled conclusion, and we have, with reservations, taken them to be solitary as stated in the literature.

In primates, the phylogenetic signal for group size is ∼0 [Kamilar & Cooper 2013], indicating that there are no grounds for using phylogenetic methods since it is necessary for both variables to have a correlated phylogenetic signal if autocorrelation is likely to bias the data. Table 1 gives estimates of the phylogenetic signal in the Carnivoran and Insectivoran group size and neocortex ratio data. While the signal for Insectivorans is strong, that for Carnivorans is weak-to-nonsignificant (though stronger than for primates), especially for neocortex size. Insectivoran phylogeny is, at best, poorly known, and probably reflects a simple switch between solitary and social species (*sensu* [Pérez-Barbería et al. 2007]). Insectivorans are now thought to be polyphyletic, having been something of a dumping ground for anything small-bodied that did not fit in any other taxonomic group. Nonetheless, current efforts to understand their taxonomy distinguish between two main genetic clades: the Eulipotyphla (true insectivores, a more primitive lineage distantly related to the ungulates, carnivores and bats) and the Afrosoricida (a more derived group of largely African origin related to hyrax, elephants and sea cows) [Beck et al 2006; Springer 2022]. To minimise any impact of phylogenetic autocorrelation, the main analyses are executed at genus level.

**Table 1.** Phylogenetic signal in Carnivoran and Insectivoran data.

| | Trait | Pagel $\lambda$ | $p(\lambda)$ | Blomberg K | $p(K)$ |
| --- | --- | --- | --- | --- | --- |
| 5 | **Carnivorans** |  |  |  |  |
|  | Group size | 0.65 | 0.09 | 0.58 | 0.02 |
|  | Neocortex ratio | 0.33 | 0.48 | 0.44 | 0.16 |
|  | Log<sub>10</sub> Group size | 0.85 | 0.04 | 0.62 | 0.01 |
|  | Log<sub>10</sub> Neocortex ratio | 0.50 | 0.24 | 0.51 | 0.04 |
| 10 | **Insectivorans** |  |  |  |  |
|  | Group size | 1.00 | 0.0001 | 0.86 | 0.005 |
|  | Neocortex ratio | 1.00 | 0.0003 | 0.83 | 0.001 |
|  | Log<sub>10</sub> Group size | 1.00 | 0.0002 | 0.81 | 0.10 |
| 15 | Log<sub>10</sub> Neocortex ratio | 1.00 | 0.0002 | 0.94 | 0.001 |

The data are provided in online *SI Dataset*. Statistical tests are 1-tailed where we test directional hypotheses.

## Results

Fig. 1 plots mean genus group size against mean genus neocortex ratio for Carnivorans and Insectivorans, superimposed on the grade lines for primates from [Dunbar & Shultz 2021a] (shown as narrow regression lines: the data for these are shown in Fig. S2]. The dashed regression line (V) gives the internal substructure layer observed in primates (see [Dunbar & Shultz 2021a]). The dotted lines are the RMA regressions for the two Insectivoran taxa. The equivalent results at species level are shown in Fig. S3. The regression lines for the individual Insectivoran and Carnivoran families are shown in Fig. S4.

**Fig. 1.**
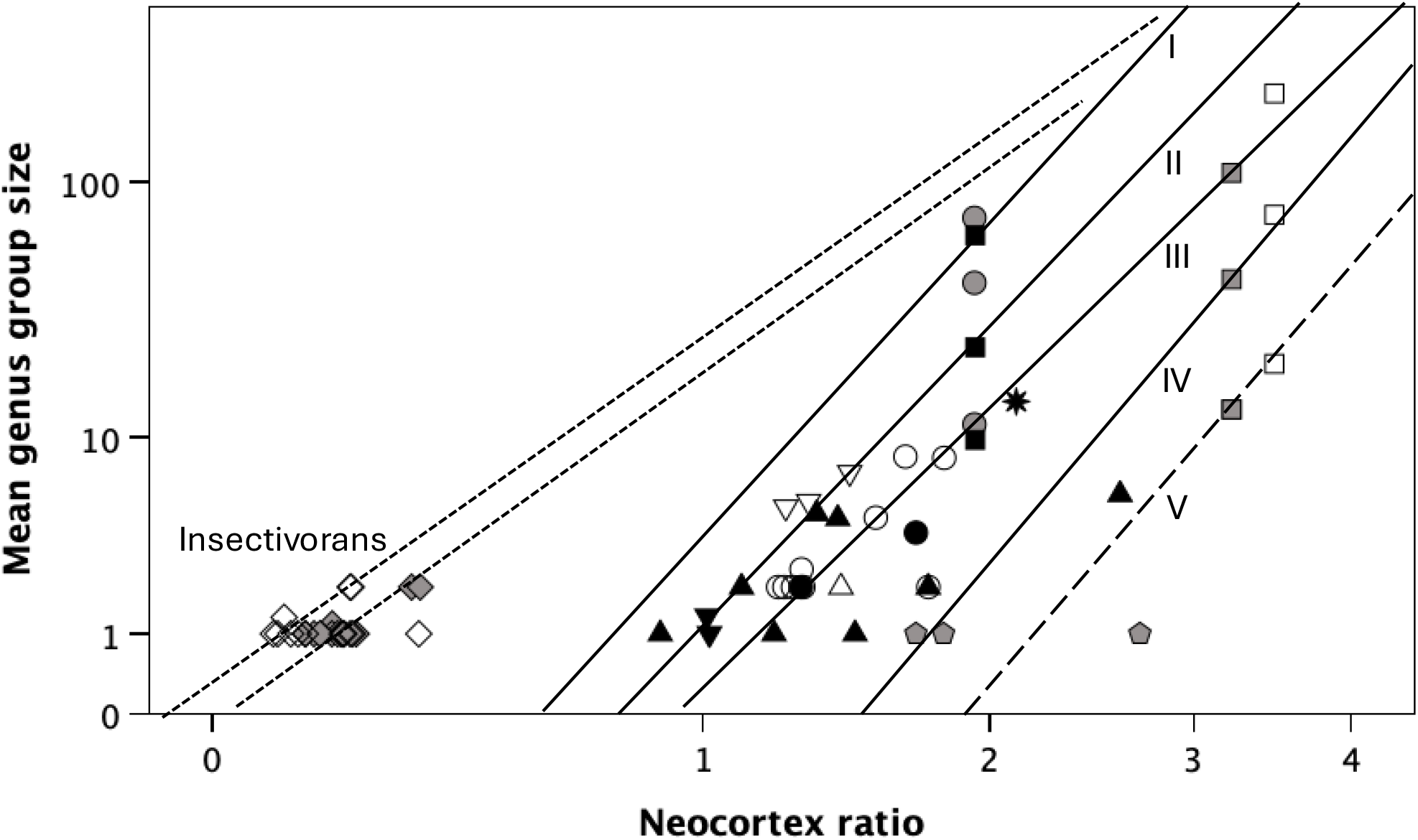
Mean genus group size for the two major Insectivoran taxa (Eulipotyphla and Afrosoricida) and nine families of carnivores plotted against mean genus neocortex ratio. In the case of spotted hyaena (*Crocuta*), separate datapoints are given for the three layers identified in their social arrangements (clans, multiclans and mega-clans). Also plotted on the graph are the mean social layer sizes for elephants (family group, bond group, clan) and orcas (pod, clan, community), along with the equivalent values for chimpanzees (clique, community, mega-community). The four grades in the primate social brain relationship (identified by Roman numerals I-IV) are shown as fine solid lines; lower dashed line identifies the internal substructure layer for primate species that have multilevel social systems (V); the upper dotted lines identify the RMA regressions set through the Eulipotyphla and Afrosoricid Insectivorans, respectively. • hyaenids; ○ spotted hyaena, *Crocuta* (top to bottom: mega-clan, multiclan, clan); ○ canids; ▴ mustelids; △ red panda; ▽ mephitids (skunks) ; ▿ procyonids (raccoons); * lion; 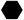 [grey pentangle]: bears; ◊ Afrosoricida Insectivorans; ◊ Eulipotyphla Insectivorans; ◼ elephants (top to bottom: clan, bond group, family group); □ chimpanzee (top to bottom: mega-community, community, clique); □ orca (top to bottom: community, clan, pod).

There is a clear distinction between the two Insectivoran taxa and the Carnivorans. Both Insectivoran groups lie well to the left of the primate distribution (comparison of intercepts with the most leftward of the primate grades: Afrosoricids: t_13_=-29.14, p<0.0001; Eulipotyphlids: t_11_=-25.05, p<0.0001). In contrast, the Carnivorans as a group map onto the primate distribution rather closely, notwithstanding the fact that a small number of individual genera lie beyond the primate range.

The OLS regressions for individual taxonomic families for which more than two genera were sampled, and for the four primate grades, are given in Table 2. Notwithstanding the statistical issues noted above, the overall OLS regression for the Carnivorans as a whole is highly significant (p<0.0001). Aside from the ursids, all the family-level regressions are positive, and significant or near significant. Taken together, this set of individual regression slopes is significantly more positive than would be expected by chance if there was no underlying relationship (Fisher meta-analysis: χ^2^=23.25, df = 2*4 families = 8, p=0.003). Although the slopes for the Carnivoran genera are more variable than those for the primate grades, they do not, as a set, differ significantly in slope (mean slopes: primates = 4.014, carnivores = 3.512; t_6_=0.32, p=0.758 2-tailed).

**Table 2.** Regression analysis for log_10_GroupSize regressed on log_10_NeocortexRatio for individual taxa, for genus-level data.

|  |  |  |  |  |  |  |  |
| --- | --- | --- | --- | --- | --- | --- | --- |
| 5 | Regression statistics |  |  |  |  |  |  |
|  | Family | grade | slope | r<sup>2</sup> | F | df | p* |
| 10 | Primates: | I | 3.668 | 0.865 | 44.75 | 1,7 | 0.0003 |
|  |  | II | 3.303 | 0.939 | 122.46 | 1,8 | <<0.0001 |
|  |  | III | 4.014 | 0.906 | 125.52 | 1,13 | <<0.0001 |
|  |  | IV | 5.072 | 0.995 | 181.52 | 1,1 | <<0.0001 |
| 15 | Eulipotyphla |  | 0.840 | 0.721 | 28.40 | 1,11 | 0.0002 |
|  | Afrosoricida |  | 0.192 | 0.063 | 0.68 | 1,10 | 0.214 |
|  | All carnivores |  | 0.477 | 0.358 | 30.17 | 1,54 | <<0.0001 |
|  | Hyaenidae† | III | 7.647 | 0.729 | 2.68 | 1,1 | 0.175 |
|  | Canidae | III | 2.768 | 0.514 | 7.59 | 1,7 | 0.015 |
|  | Mustelidae | II/(III) | 1.471 | 0.386 | 3.78 | 1,6 | 0.050 |
|  | Procyonidae (raccoons) | II | 2.309 | 0.955 | 21.40 | 1,1 | 0.068 |
| 20 | Ursidae | ?III/IV |  |  |  |  |  |
|  | Mephitidae (skunks) | II |  |  |  |  |  |
|  | *Panthera* (lion) | III |  |  |  |  |  |
|  | *Ailurus* (red panda) | III |  |  |  |  |  |
|  | * 1-tailed probability of a positive slope [negative slopes disprove the hypothesis] |  |  |  |  |  |  |
| 25 | † using only basal clan size |  |  |  |  |  |  |

Although both Insectivoran taxa exhibit a positive correlation between group size and brain size (Fisher’s meta-analysis: χ^2^=20.12, df = 4, p=0.0005), in fact this is mainly due to a simple monogamy effect: pairbonded genera have significantly larger neocortex ratios than solitary genera (Eulipotyphlids: neocortex volume, t_10_=2.31, p=0.022; neocortex ratio, t_10_=7.25, p<0.001; Afrosoricids: neocortex volume, t_9_=1.53, p=0.081; neocortex ratio, t_9_=0.77, p=0.230). Carnivorans exhibit a similar effect: although pairbonded genera (those with a mean group size of N<5) do not have significantly larger neocortex ratios than solitary genera (t_19_=0.77, p=0.226 1-tailed), they do have significantly larger neocortices (t_19_=1.86, p=0.039 1-tailed).

Note how closely the datapoints for elephants, orcas and chimpanzees sit to the primate grade lines. This is also true for two of the three *Crocuta* datapoints: their basal clan size (essentially a matriline) and the larger of the two multi-clan units fit closely to grades I and III; however, the intermediate level group (the smaller multiclan grouping) sits between grades I and II, suggesting that do not form a formal fractal series in the way the others do.

Two other points should be noted. First, a small number of carnivore genera lie to the right of the primate distribution and appear to be exceptional even for the other members of their families. These include *Pteronura* (giant otter), *Eir*a (tayra), *Taxidea* (American badger), *Chrysocyon* (maned wolf) and the ursids as a group. All of these seem to have unusually large neocortex ratios for their group sizes (which in most cases are described as solitary or pairbonded). Second, several of the Insectivorans (but especially the moles and the giant otter shrew, *Potamogale*) lie to the right of all the other Insectivorans (of both clades). Again, these are described as being solitary, but have neocortex ratios commensurate with pairliving.

## Discussion

These analyses indicate that, despite a number of claims to the contrary, Insectivorans, Carnivorans, elephants and orcas all fall in line with the primate social brain relationship. Insectivorans differ from the other taxa in that they lie on a separate grade (or pair of grades) to the left the primate social brain graph. This suggests that Insectivoran sociality is likely to be of a different quality to that observed in the other mammals in that it does not require significant computational or processing power. One reflection of this may be that their form of monogamy lacks the intensity of pairbonding found in anthropoid primates, canids and some bovids. The Carnivorans, in contrast, map closely onto the primate range. While there is certainly a strong solitary-versus-pairbonded effect (as noted by [Perez-Barberia et al. 2007; Shultz & Dunbar 2007]), there is a linear effect identical to that seen in primates mapped on top of this for a small number of genera that live in social groups >5 (just 12 genera out of the 28 sampled). The monogamy effect is consistent with the claim that sociality initially arose via some form of pairliving[Kappeler & Pozzi 2019]. Unlike the primates, it is only within some Carnivoran lineages that a quantitative signal is found, although even then it spans only a very modest part of the extended range we find in primates.

In an attempt to explain their own negative results, Finarelli & Flynn [2011] asserted that previous analyses for Carnivorans had relied almost exclusively on canids. In fact, this claim was untrue then and is untrue in the present case: the current analysis uses the same data as the earlier analyses and includes all major Carnivoran families (although the felids are represented by only one species, the lion). It is notable that the lion sits very squarely on grade III with the smarter primates. Shultz & Dunbar [2010], who included data on all felids, noted that the felids are unique among the major mammalian sub-orders in exhibiting no significant change in relative brain size across geological time (see their Fig. 1), a finding commensurate with the fact that all felid genera except the lion and cheetah are solitary.

Inspection of Fig. 1 suggests that the Carnivoran genera divide between two specific primate grades: skunks (Mephitidae), raccoons (Procyonidae) and, with a few notable exceptions, the mustelids sit with the less social monkeys on or near primate grade II, while the hyaenids, canids, lion and red panda sit on grade III with the more social monkeys. Of the 30 Carnivoran genera in the sample, only one canid, three mustelids and the three ursids (23% of genera) lie outside the 95% CIs for the two inner primate grade lines (grades II and III). The parallelogram defined by the outer CIs for these two grades (see Fig. S4) occupies 48.7% of the state space within which Carnivorans occur. The probability of ≤7/30 datapoints lying outside these limits if they were randomly distributed in the state space is p=0.0017 (2-tailed).

The non-ursid exceptions (all of which are monospecific genera) include the maned wolf (*Chrysocyon*), the giant otter (*Pteronura*), tayra (*Eira*) and American badger (*Taxidea*), all of whom sit on or near primate grade IV (i.e. with the apes). It is not obvious why these genera should be positioned so far from their congenors, since it implies that they have much larger neocortices for their observed group sizes than would be expected. There seems to be nothing in either their social skills or their cognitive capacities to justify such large neocortices, though none are well studied. Given the poor knowledge of their natural history, however, it is possible that their group sizes are underestimated. If these genera sat on grade III with the other mustelids and canids, *Taxidea* would have a predicted group size of 3.2 (essentially a monogamous pair with offspring), while *Eira* and the maned wolf would have predicted group sizes of ∼6.5 (similar to wolves). This is not entirely implausible. Less easy to explain is *Protonura* (giant otter): its neocortex ratio would predict a mean group size of ∼30 if it sat on grade III with the rest of the mustelids. This seems implausibly large, although otters in general are extremely social and *Protonura*, in particular, has been described as intensely social, strongly pairbonded with a large, frequent physical contact including allogrooming, a complex vocal repertoire and living in very stable groups up to 20 individuals [Duplaix 1980].

The same conclusion seems to apply to the Insectivoran exceptions. If the giant otter shrew was on the Afrosoricid line, it would have a predicted group size of 3.5; *Limnogale* (one of several tenrec genera), *Chrysochloris* (the golden mole) and *Potamogale* and *Micropotamogale* (the small otter shrews) would all have predicted group sizes of 2. This would put all of them in the pairbonded monogamy category along with the other large-brained genera. At least some of these taxa live in overlapping territories, even though all forage alone [Kingdon 2014]. More detailed study of their natural history may reveal that individuals share nests and/or have positive affiliative relationships with a preferred partner.

The bear family as a whole is the real puzzle. Two of the three genera sampled fall close to grade IV (the grade associated with the apes), but the sun bear (*Helarctos*) falls far beyond even this grade and stands out as anomalous. The claim that bears are on a cognitive par with apes seems extravagant. It is possible, however, that we misjudge the social world of these species – yet another case where the foraging group has been mistaken for the social group. If the bears actually fell on grade III (which might not be unreasonable given their manipulative and foraging skills), then black bears (*Ursus*) and the giant panda (*Ailuropoda*) would be predicted to live in communities that numbered ∼6.5 individuals – not large by primate standards, but perfectly feasible as a small orangutan-like dispersed community where individuals range in a common territory and have broadly affiliative relationships with each other (and agonistic ones with neighbours). While it is unlikely that the brains of all three genera have been overestimated, it is possible that, like the dolphin [Hof, Chanis & Marino, 2005; Marino *et al*., 2007; Oelschläger, 2008], their brains are organised differently, with large neocortical regions devoted to an ecologically crucial function (e.g. managing a heavy body mass clambering about in trees). Once again, however, we need to ask whether there is something about ursid sociality and/or cognition that we do not yet understand.

The fact that the two of the three grouping sizes for the spotted hyaena (the basal matrilineal clan and the mega-clan comprising many basal clans [Dunbar 2024b]) sit on different grades (II and III) is particularly interesting. It parallels a similar finding for a number of primate genera (including *Miopithecus, Chlorocebus, Piliocolobus, Papio hamadryas* and *Theropithecus*) that have multilevel social systems based on fission-fusion principles [Dunbar & Shultz a]. Hyaenids share with grade III/IV primates sophisticated forms of social cognition (notably reconciliation and third party recognition [Wahaj et al. 2001; Engh et al. 2005]) that play an important role in primates’ especially advanced form of sociality [Dunbar 2025]. In all these cases, the larger grouping is achieved by deferring group fission, such that what would otherwise have been two daughter groups remain together as a weakly bonded supergroup when ecological circumstances make this desirable (and possible) [Dunbar 2025]. The fact that the intermediate grouping level (the multiclan) does not sit on grade II is significant, because it suggests that this species does not have a form of fractally structured multilevel sociality like that found in anthropoid primates [Dunbar & Shultz 2021a], elephants and orcas (Fig. 1).

It seems that the reason why some (but not all) previous analyses failed to find a social brain relationship is that they used extracranial volumes rather than neocortex size (the actual basis for the social brain hypothesis) as a measure of brain size, combined with the fact that they ignored the grades present in the distribution of these data (resulting in lowered regression slopes due to Simpson’s Paradox). The present findings indicate that statistically better informed analyses yield rather more convincing and interesting results.

The fact that the partition between less and more social Carnivoran genera map onto the primate grades II and III prompts questions about the cognitive abilities of these taxa, especially in the social domain. The issue is whether Carnivorans have evolved different forms of cognition (in the way dolphins have evolved echolocation) or whether the demands of sociality in the two orders have resulted in convergence on similar forms of social cognition. With the exception of the spotted hyaena [Johnson-Ulrich & Holekamp 2020] and two canids [MacLean et al. 2014], both of whom sit on grade III (Fig. 1) with grade III-like cognitive competences, relevant data are not widely available for Carnivorans. Efforts are clearly needed to assess a wider range of carnivore species on standard cognitive tasks, both to compare the less and more social taxa and to compare these with their primate equivalents.

In sum, contrary to previous assumptions, both the Insectivorans and Carnivorans comply with the social brain hypothesis in its fully quantitative form, even if they lie at the lower ends of the group size distribution on these grades, and hence do not live in such large groups as primates do. Since primate social evolution is now known to depend on high order socio-cognitive abilities needed to counteract the very powerful centrifugal forces that drive animals apart, in particular mentalising (understanding intentions) and the capacity for self- control [Dunbar & Shultz 2021b, 2025; Dunbar 2025], this raises fundamental questions about how other mammalian orders compare with primates on these abilities and whether it is these capacities or, more simply, the structural features of network size as attractors [West et al. 2023] that result in the similarities between the primates and other mammalian orders demonstrated here.

## References

Avelino-de-Souza, K., Mynssen, H., Chaim, K., Parks, A. N., Ikeda, J. M., Cunha, H. A., … & Patzke, N. (2024). Anatomical and volumetric description of the Guiana dolphin (Sotalia guianensis) brain from an ultra-high-field magnetic resonance imaging. Brain Structure and Function 229: 1889–1911.

Beck, R. M., Bininda-Emonds, O. R., Cardillo, M., Liu, F. G. R. & Purvis, A. (2006). A higher-level MRP supertree of placental mammals. BMC Evolutionary Biology 6: 1–14.

Chambers, H. R., Heldstab, S. A. & O’Hara, S. J. (2021). Why big brains? A comparison of models for both primate and carnivore brain size evolution. PLoS One 16: e0261185.

Dheer, A. (2016). Resource partitioning between spotted hyenas (Crocuta crocuta) and lions (Panthera leo). MSc thesis, University of Southampton.

Dunbar, R.I.M. (1993). Coevolution of neocortex size, group size and language in humans. Behavioral and Brain Sciences 16: 681–735.

Dunbar, R.I.M. (2024). Multilevel sociality in the spotted hyaena: how to live in large groups without falling prey to the infertility trap. African Journal of Ecology 62: e13277.

Dunbar, R.I.M. (2025). Structural and cognitive mechanisms of group cohesion in primates. Behavioral and Brain Sciences (in press).

Dunbar, R.I.M. & Bever, J. (1998). Neocortex size predicts group size in carnivores and some insectivores. Ethology 104: 695–708.

Dunbar, R.I.M. & Shultz, S. (2021a). Social complexity and the fractal structure of social groups in primate social evolution. Biological Reviews 96: 1889–1906.

Dunbar, R.I.M. & Shultz, S. (2021b). The infertility trap: the fertility costs of group-living in mammalian social evolution. Frontiers in Ecology and Evolution 9: 634664.

Dunbar, R.I.M. & Shultz, S. (2023). Four errors and a fallacy: pitfalls for the unwary in comparative brain analyses. Biological Reviews 98: 1278–1309.

Dunbar, R.I.M. & Shultz, S. (2025). The social role of self-control. BioRxiv /2024/578865.

Dunbar, R.I.M., MacCarron, P. & Shultz, S. (2018). Primate social group sizes exhibit a regular scaling pattern with natural attractors. Biology Letters 14: 20170490.

Duplaix, N. (1980). Observations on the ecology and behavior of the giant river otter Pteronura brasiliensis in Suriname. Revue d’Écologie 34: 495–620.

Engh, A. L., Siebert, E. R., Greenberg, D. A., & Holekamp, K. E. (2005). Patterns of alliance formation and postconflict aggression indicate spotted hyaenas recognize third-party relationships. Animal Behaviour 69: 209–217.

Finarelli, J. A. & Flynn, J. J. (2009). Brain-size evolution and sociality in Carnivora. Proceedings of the National Academy of Sciences, USA, 106: 9345–9349.

Frank, L. G. (1986). Social organization of the spotted hyaena (Crocuta crocuta). I. Demography. Animal Behaviour 34: 1500–1509.

Gittleman, J. L. (1986). Carnivore brain size, behavioral ecology, and phylogeny. Journal of Mammalogy 67: 23–36.

Haug, H. (1970). Quantitative data in neuroanatomy. Progress in Brain Research 33: 113–127.

Hakeem, A. Y., Hof, P. R., Sherwood, C. C., Switzer III, R. C., Rasmussen, L. E. L., & Allman, J. M. (2005). Brain of the African elephant (Loxodonta africana): neuroanatomy from magnetic resonance images. Anatomical Record Part A 287: 1117–1127.

Henschel, J. R. (1986). The socio-ecology of a spotted hyaena Crocuta crocuta clan in the Kruger National Park. PhD thesis, University of Pretoria

Hill, R.A., Bentley, A. & Dunbar, R.I.M. (2008). Network scaling reveals consistent fractal pattern in hierarchical mammalian societies. Biology Letters 4: 748–751.

Hof, P. R., Chanis, R. & Marino, L. (2005). Cortical complexity in cetacean brains. Anatomical Record A 287, 1142–1152.

Holekamp, K. E., & Dloniak, S. M. (2010). Intraspecific variation in the behavioral ecology of a tropical carnivore, the spotted hyena. In Advances in the Study of Behavior 42: 189–229.

Holekamp, K. E., Sakai, S. T. & Lundrigan, B. L. (2007). Social intelligence in the spotted hyena (Crocuta crocuta). Philosophical Transactions of the Royal Society, London, 362B: 523–538.

Holekamp, K. E., Cooper, S. M., Katona, C. I., Berry, N. A., Frank, L. G., & Smale, L. (1997). Patterns of association among female spotted hyenas (Crocuta crocuta). Journal of Mammalogy 78: 55–64.

Johnson-Ulrich, L. & Holekamp, K. E. (2020). Group size and social rank predict inhibitory control in spotted hyaenas. Animal Behaviour 160: 157–168.

Kamilar, J.M. & Cooper, N. (2013). Phylogenetic signal in primate behaviour, ecology and life history. Philosophical Transactions of the Royal Society, London, 368B: 20120341.

Kamiya, B. T. & Pirlot, P. (1988). The brain of the lesser panda Ailurus fulgens: A quantitative approach. Journal of Zoological Systematics and Evolutionary Research 26: 65–72.

Kamtya, T. & Pirlot, P. (1988). The brain of the Malayan bear (Helarctos malayanus). Journal of Zoological Systematics and Evolutionary Research 26: 225–235.

Kappeler, P. M. & Pozzi, L. (2019). Evolutionary transitions toward pair living in nonhuman primates as stepping stones toward more complex societies. Science Advances 5: eaay1276.

Kingdon, J. (2014). Mammals of Africa: Volume I: Introductory Chapters and Afrotheria. A&C Black.

Kruuk, H. (1966). Clan-system and feeding habits of spotted hyaenas (Crocuta crocuta Erxleben). Nature 209: 1257–1258.

Marino, L., Connor, R. C., Fordyce, R. E., Herman, L. M., Hof, P. R., Lefebvre, L., Lusseau, D., McGowan, B., Nimchinsky, E. A., Pack, A. A., Rendell, L., Reidenberg, J. S., Reiss, D., Uhren, M. D., Van der Guht, E. & Whitehead, H. (2007). Cetaceans have complex brains for complex cognition. PLoS Biology 5, e139.

Mars, R. B., Neubert, F.-X., Noonan, M. P., Sallet, J., Toni, L. & Rushworth, M. F. S. (2012). On the relationship between the “default mode network” and the “social brain”. Frontiers in Human Neuroscience 6: 189.

Müller, A. E. & Thalmann, U. R. S. (2000). Origin and evolution of primate social organisation: a reconstruction. Biological Reviews 75: 405–435.

Nekaris, A. & Bearder, S. K. (2007). The Lorisiform primates of Asia and mainland Africa. In: C. Campbell, A. Fuentes, K. Mackinnon, S. Bearder & J. Stumpf (eds.) Primates in Perspective, pp. 24–45. Oxford: Oxford University Press.

Oelschläger, H. H. (2008). The dolphin brain—a challenge for synthetic neurobiology. Brain Research Bulletin 75, 450–459.

Pérez-Barbería, J., Shultz, S. & Dunbar, R.I.M. (2007). Evidence for intense coevolution of sociality and brain size in three orders of mammals. Evolution 61: 2811–2821.

Rademacher, J., Morosan, P., Schormann, T., Schleicher, A., Werner, C., Freund, H. J., & Zilles, K. (2001). Probabilistic mapping and volume measurement of human primary auditory cortex. Neuroimage 13: 669–683.

Ridgway & Au (2009). Hearing and echolocation in dolphins. Encyclopedia of Neuroscience 4: 1031–109. Elsevier.

Röhrs, V. M. (1986). Cephalisation, Telencephalisation und Neocorticalisation bei Mustelidae. Journal of Zoological Systematics and Evolutionary Research 24: 157–166.

Röhrs, V. M. (1986). Cephalisation bei Caniden. Journal of Zoological Systematics and Evolutionary Research, 24(4), 300–307.

Röhrs, V. M., Ebinger, P. & Weidemann, W. (1989). Cephalisation bei Viverridae, Hyaenidae, Procyonidae und Ursidae 1. Journal of Zoological Systematics and Evolutionary Research 27: 169–180.

Rushworth, M. F., Mars, R. B. & Sallet, J. (2013). Are there specialized circuits for social cognition and are they unique to humans? Current Opinion in Neurobiology 23: 436–442.

Sakai, S. T., Arsznov, B. M., Lundrigan, B. L. & Holekamp, K. E. (2011). Brain size and social complexity: a computed tomography study in hyaenidae. Brain Behavior and Evolution 77: 91–104.

Shultz, S. & Dunbar, R.I.M. (2007). The evolution of the social brain: Anthropoid primates contrast with other vertebrates. Proceedings of the Royal Society, London, 274B: 2429–2436.

Shultz, S. & Dunbar, R.I.M. (2010). Encephalisation is not a universal macroevolutionary phenomenon in mammals but is associated with sociality. Proceedings of the National Academy of Sciences, USA, 107: 21582–21586.

Shultz, S. & Dunbar, R.I.M. (2022). Socioecological complexity in primate groups and its cognitive correlates. Philosophical Transactions of the Royal Society, London, 377B: 20210296.

Smith, J. E., Kolowski, J. M., Graham, K. E., Dawes, S. E. & Holekamp, K. E. (2008). Social and ecological determinants of fission–fusion dynamics in the spotted hyaena. Animal Behaviour 76: 619–636.

Springer, M. S. (2022). Afrotheria. Current Biology 32: R205–R210.

Stephan, H., Frahm, H. & Baron, G. (1981). New and revised data on volumes of brain structures in insectivores and primates. Folia Primatologica 35: 1–29.

Tilson, R. L., & Henschel, J. R. (1986). Spatial arrangement of spotted hyaena groups in a desert environment, Namibia. African Journal of Ecology 24: 173–180.

Vullioud, C., Davidian, E., Wachter, B., Rousset, F., Courtiol, A., & Höner, O. P. (2019). Social support drives female dominance in the spotted hyaena. Nature Ecology & Evolution 3: 71–76.

Wahaj, S. A., Guse, K. R., & Holekamp, K. E. (2001). Reconciliation in the spotted hyena (Crocuta crocuta). Ethology 107: 1057–1074.

West, B., Massari, G.F., Culbreth, G., Failla, R., Bologna, M., Dunbar, R.I.M. & Grigolini, P. (2020). Relating size and functionality in human social networks through complexity. Proceedings of the National Academy of Sciences, USA, 117: 18355–18358.

West, B., Dunbar, R.I.M., Culbreth, G. & Grigolini, P. (2023). Fractal structure of human and primate social networks optimizes information flow. Proceedings of the Royal Society, London, 479A: 20230028.

Wright, A., Scadeng, M., Stec, D., Dubowitz, R., Ridgway, S., & Leger, J. S. (2017). Neuroanatomy of the killer whale (Orcinus orca): a magnetic resonance imaging investigation of structure with insights on function and evolution. Brain Structure and Function 222: 417–436.

