## Supplementary Methods for "Insectivorans and Carnivorans Exhibit the Same Social Brain Relationship as Primates"

**R.I.M. Dunbar**

**Susanne Shultz**

### Supplementary Information

The distribution of group (clan) sizes in the spotted hyaena (*Crocuta crocuta*) exhibits a distinctively multimodal pattern (Fig. S1). *k*-means cluster analysis indicates an optimal partition into three subsets with mean values at 11.3, 41.0 and 72.8 (Dunbar 2024).

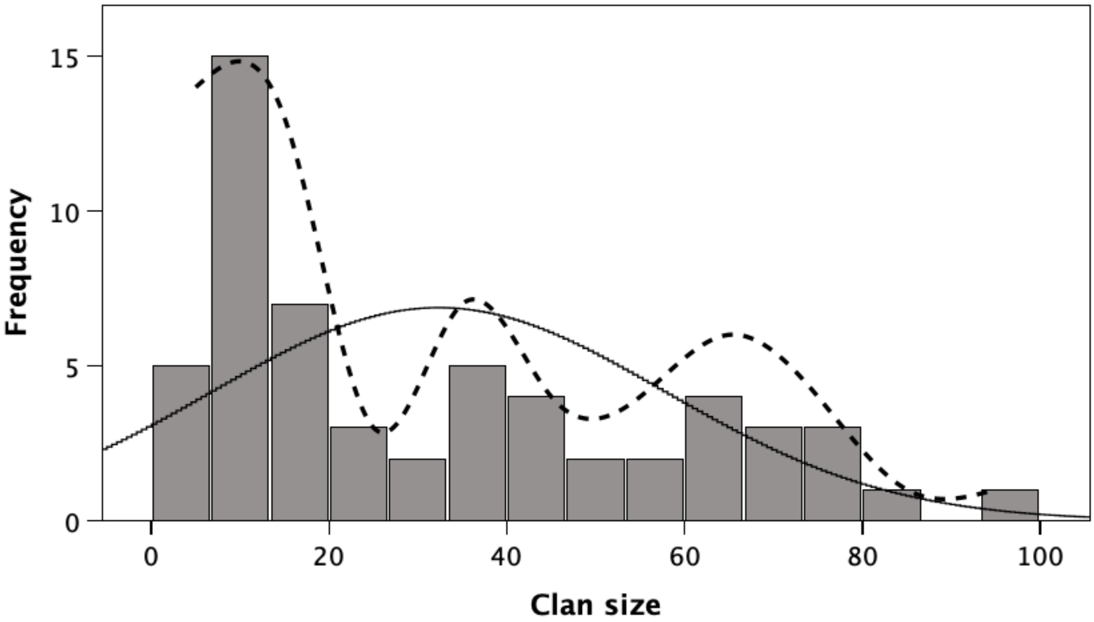


Fig. S1

Distribution of clan sizes in spotted hyaena has a trimodal form. The solid line is the normal distribution fitted to the data; the dashed line is the best-fit polynomial. Reproduced from Dunbar (2024).

Fig. S2 plots the original data for primates to illustrate the natural grades in the social brain data. The grades were identified by *k*-means cluster analysis of the residuals from the RMA regression for the full dataset (for details,see Dunbar & Shultz 2021a).

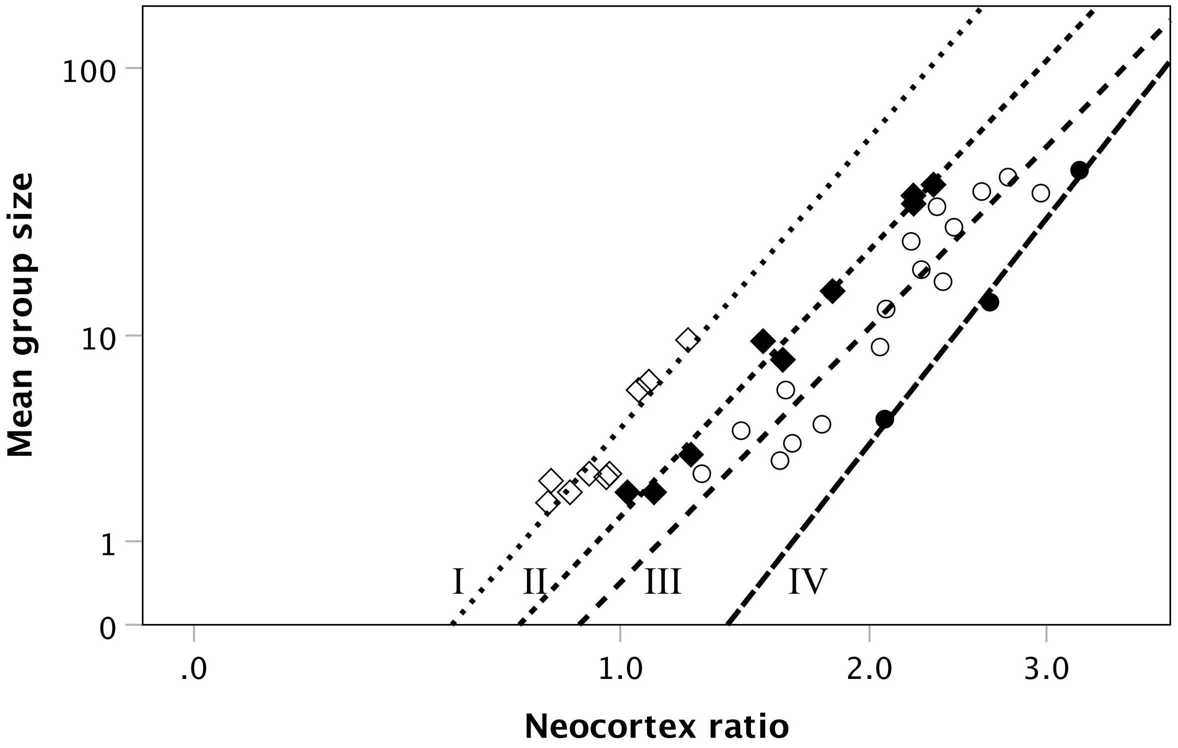


Fig. S2
Plot of primate social brain data. The four socio-economic grades are indicated by different symbols and regression lines, and identified by the Roman numerals. Grade IV on the extreme right consists only of apes.

Fig. S3 plots the data for individual species mean values.

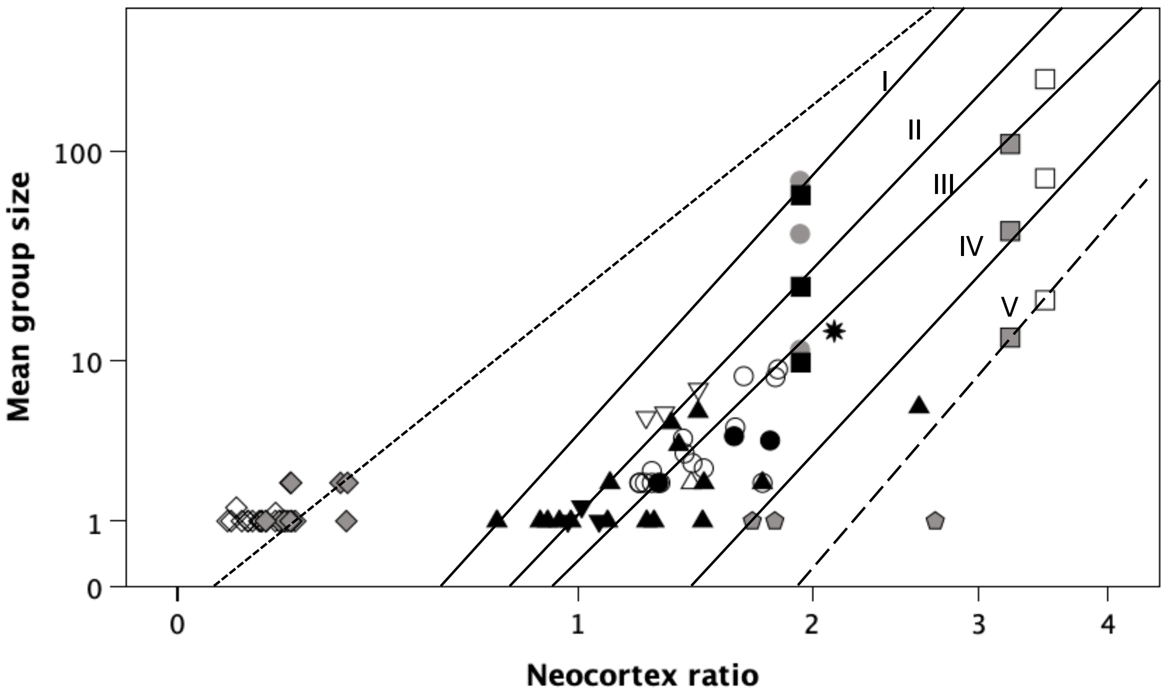


Fig. S3

Insectivoran and Carnivoran social brain data for individual species. Solid lines are the regression lines for the four primate social brain grades (I to IV); the lower dashed line is the primate grade V for the internal substructuring of groups; the upper dotted line is the RMA line for Afrosoricid Insectivorans.

● hyaenids; ○ spotted hyaena, *Crocuta* (top to bottom: mega-clan, multiclan, clan); ○ canids; ▲ mustelids; △ red panda; ▼ mephitids (skunks) ; ∇ procyonids (raccoons); ✱ lion; ⬣ [grey pentangle]: bears; ◊ Eulipotyphla Insectivorans; ◊ Afrosoricida Insectivorans; ■ elephants (top to bottom: clan, bond group, family group); □ chimpanzee (top to bottom: mega-community, community, clique); □ orca (top to bottom: community, clan, pod).

Fig. S4 plots the social brain data for individual insectivore and carnivore families in the form of genus mean values.

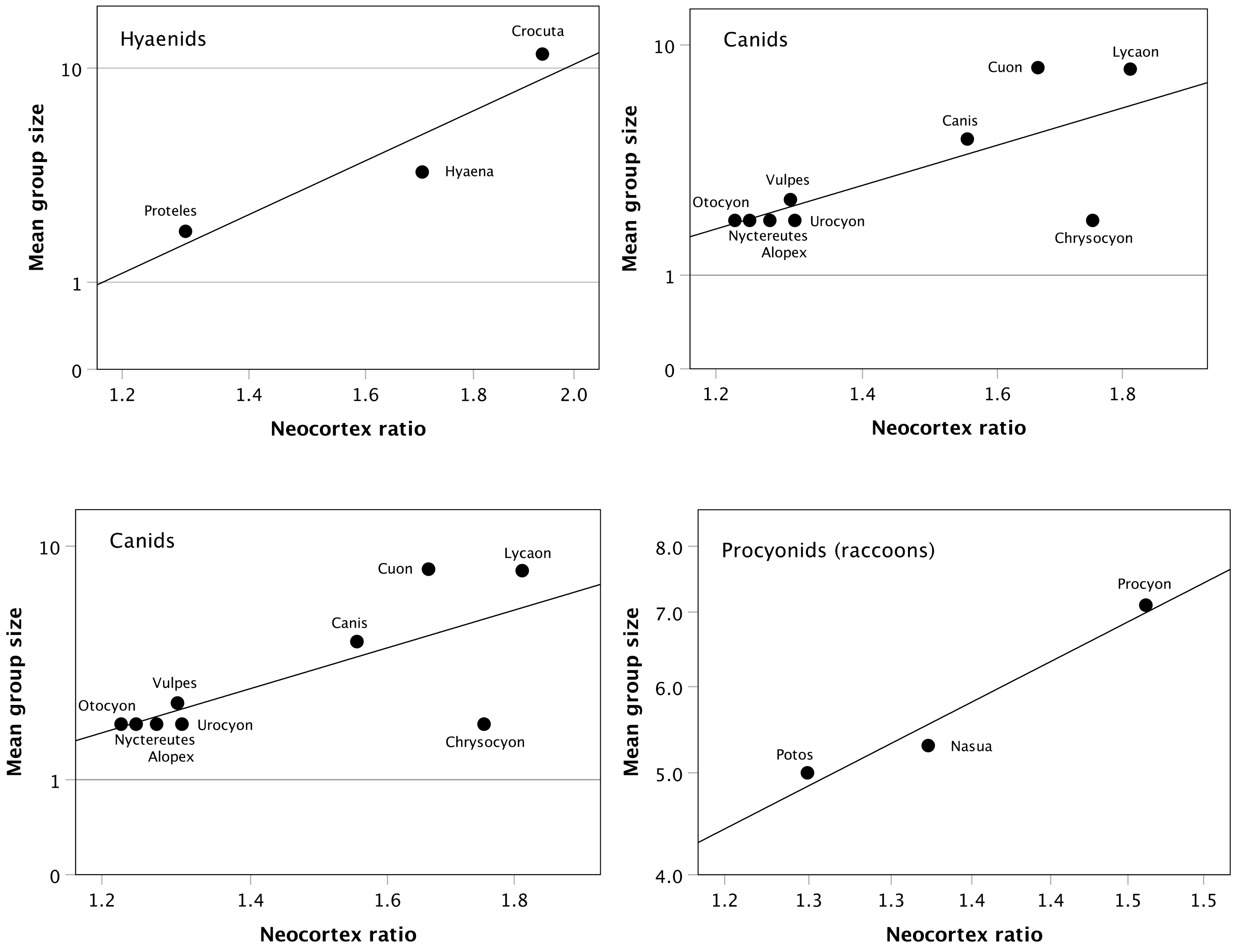

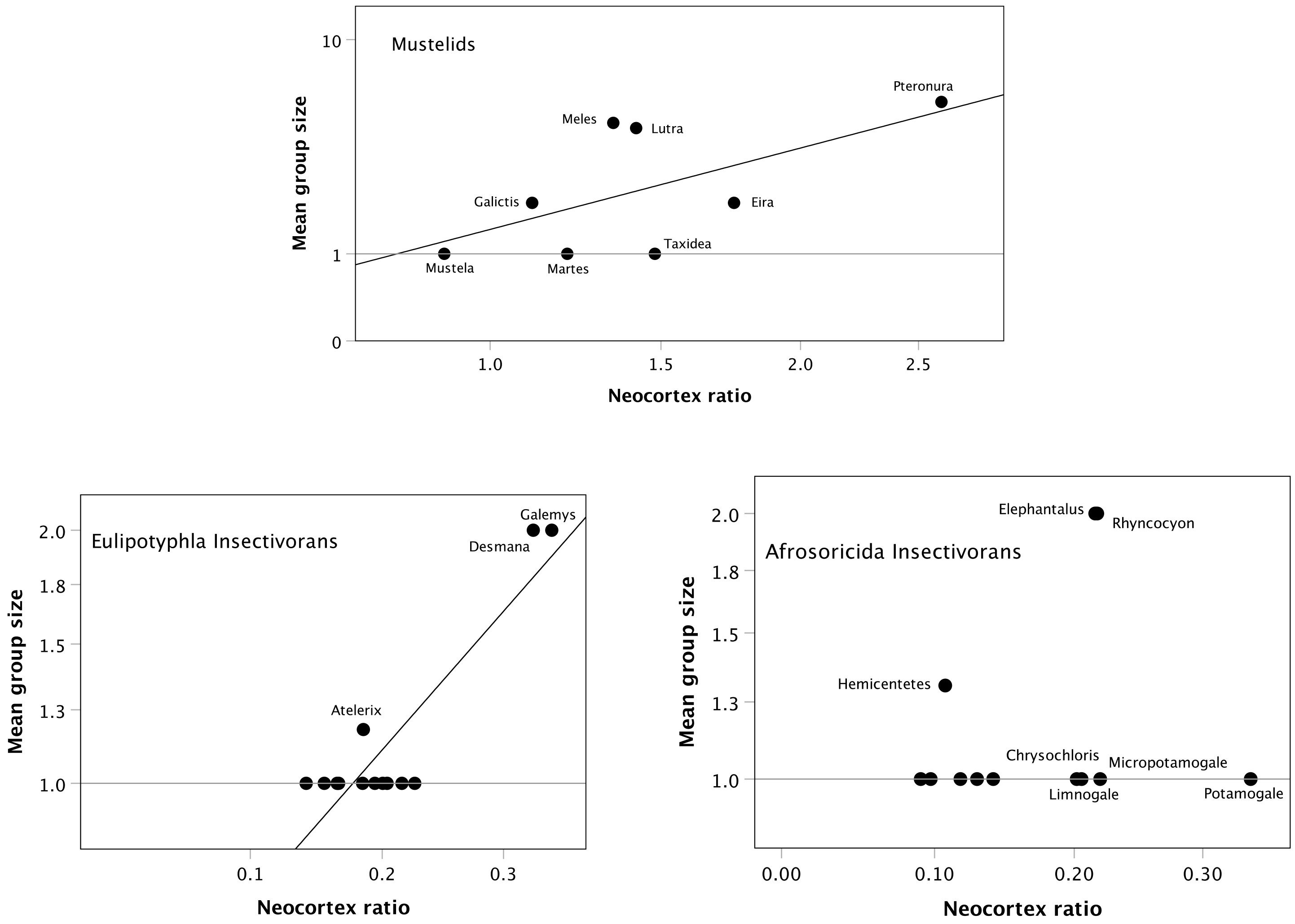


Fig. S4

Plots for individual carnivore and Insectivoran families with N>2 genera. Data are generic means.

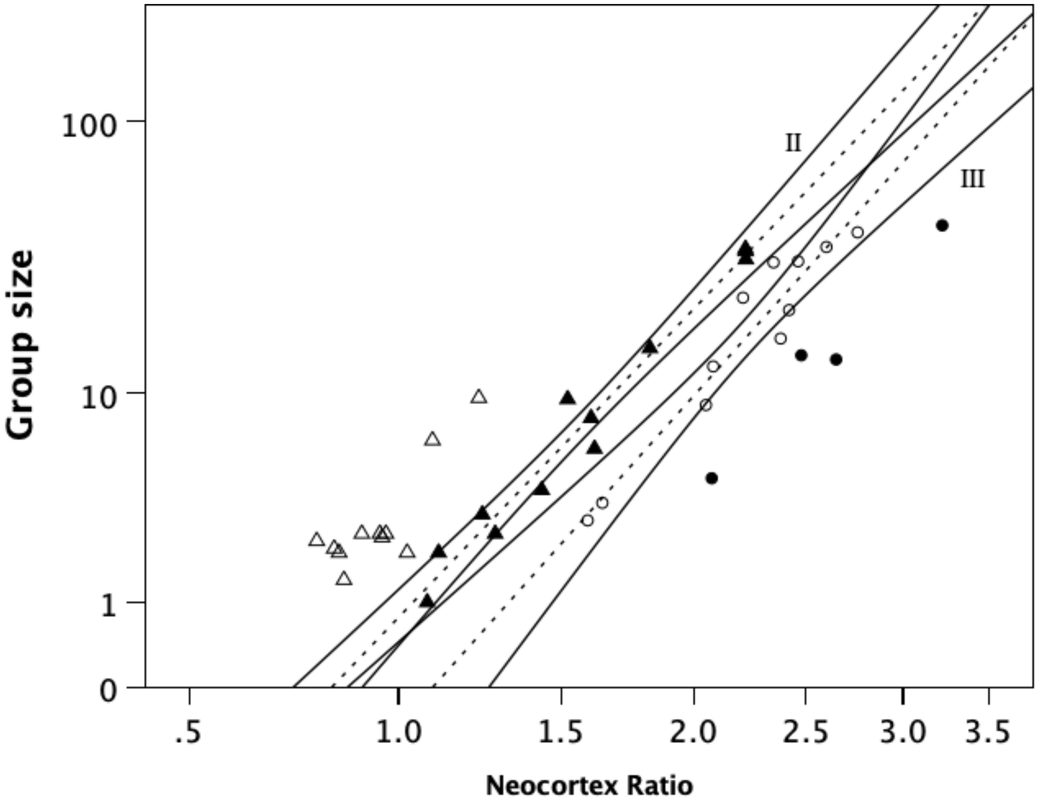


Fig. S5

Primate social brain date with 95% prediction CIs for grades II and III. Dotted lines are the OLS regressions for grades II and III; the solid lines are the 95% prediction CIs around these regressions.
